# Estimating the distribution of livestock in space and time: a time-series contribution to livestock density mapping

**DOI:** 10.1101/2022.02.20.479787

**Authors:** Julianne Meisner, Agapitus Kato, Marshall Lemerani, Erick Mwamba Miaka, Acaga Ismail Taban, Jonathan Wakefield, Ali Rowhani-Rahbar, David Pigott, Jonathan Mayer, Peter Rabinowitz

## Abstract

More than one billion people rely on livestock for income, nutrition, and social cohesion, however livestock keeping can facilitate disease transmission and contribute to climate change. While data on the distribution of livestock thus have broad utility across a range of applications, efforts to map the distribution of livestock on a large scale are limited to the Gridded Livestock of the World (GLW) project. We present a complimentary effort to map the distribution of cattle and pigs in Malawi, Uganda, Democratic Republic of Congo (DRC), and South Sudan. In contrast to GLW, which uses dasymmetric modeling applied to census data to produce time-stratified estimates of livestock counts and spatial density, our work uses complex survey data and distinct modeling methods to generate a time-series of livestock distribution, defining livestock density as the ratio of animals to humans. In addition to favorable cross-validation results and general agreement with national density estimates derived from external data on national human and livestock populations, our results demonstrate extremely good agreement with GLW-3 estimates, supporting the validity of both efforts. Our results furthermore offer a high-resolution time series result and employ a definition of density which is particularly well-suited to the study of livestock-origin zoonoses.

## Introduction

Globally, one billion people living on less than US$2 per day depend on livestock. For these communities, which include 80% of the poor in Africa, livestock are a critical source of household income, transport, draft power for crop agriculture, and nutrition, providing 11% of energy and 26% of dietary protein among the poor in East Africa, and up to 50% of energy for children under five in pastoralist communities.^1, 2^ Livestock keeping also plays an important cultural role, serving as a means to maintain family cohesion and social networks, gain political prestige, and strengthen legal claims on pasture land.^3, 4^ These critical roles livestock play, however, commonly result in close human-animal contact, driving transmission of zoonotic diseases and resulting in over 2.5 billion cases of human illness and 2.7 million deaths per year.^2^

Livestock also have major environmental impacts across a range of scales. In Africa, where an estimated 1 billion or more of the projected increase in the human population will occur, urbanization and an increased demand for animal products are also expected,^5^ driving the so-called “livestock revolution.”^6^ Production systems are generally low-input in sub-Saharan Africa, with very little technological change in the past 40 years;^5^ compounded by degraded natural resources, this lack of technological change requires increasing production demands to be met by overgrazing and land-use changes, rather than intensification, compromising biodiversity and ecosystem services.^7^ Furthermore, where intensification has been achieved, animals and animal wastes are generally concentrated, facilitating disease transmission and water pollution in the absence of proper controls.^1^

On a larger scale, livestock systems have a bi-directional relationship with climate change. Climate change affects the quantity and quality of feeds available to livestock, drives production losses due to heat stress, promotes uneven distribution of water resources, and modifies the distribution of livestock diseases and disease vectors, representing another manifestation of the disproportionate burden climate change places on the world’s resource poor. In turn, livestock and livestock systems are substantial users of natural resources, in particular water and land for grazing and feed production, and contribute to greenhouse gas emissions and climate change.^5, 8^

Thus, high spatiotemporal resolution maps of the distribution of livestock hold utility for a range of research and policy applications, from epidemiology and public health to economics and climate science. To this end, the Gridded Livestock of the World (GLW) database was developed in 2007,^9^ with GLW-3 being the most recent iteration. Published in 2018 but providing results for 2010, GLW-3 produces two sets of results, one using dasymetric modeling to disaggregate census counts, which is based on weights derived from statistical models that use spatial covariates, and a simple aerial weighting approach that produces estimates free from association with the spatial covariates used in the daysmetric results. These results are produced globally at a resolution of 0.083333° for counts of cattle, buffaloes, horses, sheep, goats, pigs, chickens, and ducks.^10^ GLW does not use any household survey data, such as the Demographic and Health Survey or the Living Standards Measurement Studies, as these data use complex survey sampling approaches that would not be well-suited to GLW’s current modeling approaches. While numerous prior authors have mapped livestock diseases and disease vectors, beyond GLW mapping of livestock itself remain isolated to highly localized efforts such as grazing intensity in Kazakhstan^11^ and livestock movements in Sahelian Africa.^12^

To build on and support GLW efforts, we have used household survey and census data on livestock ownership and the stochastic partial differential equations (SPDE) approach to Gaussian process modeling to generate high-resolution (0.017°) maps of cattle and pig density. We have produced these maps for every year from 2000-2020 for Malawi and Uganda and 2008-2015 for Democratic Republic of Congo (DRC), facilitating their use in longitudinal analyses. We have additionally used small area estimation to produce density maps at the county level for South Sudan in 2008. In all four countries, we define density as the ratio of animals to humans, producing a map in which the human population is flattened. We present here our methods and resulting products, and discuss our results in the context of GLW-3 and extension of these approaches to other countries and species.

## Results

### Input data

In the final dataset there were 6,330 clusters for cattle mapping and 6,342 clusters for pig mapping in Malawi; 3,269 clusters for cattle and 3,310 clusters for pigs in Uganda; and 861 clusters for each species in DRC. For South Sudan, data were available for all counties except for Khorflus. For the three SPDE countries (Malawi, Uganda, and DRC), cluster distribution in space is presented in Figure 1, and data richness (number of survey clusters) by year and country is summarized in Figure 2. For cattle, counts were summed over local and exotic breeds, and dairy and non-dairy herds.

**Figure 1.**
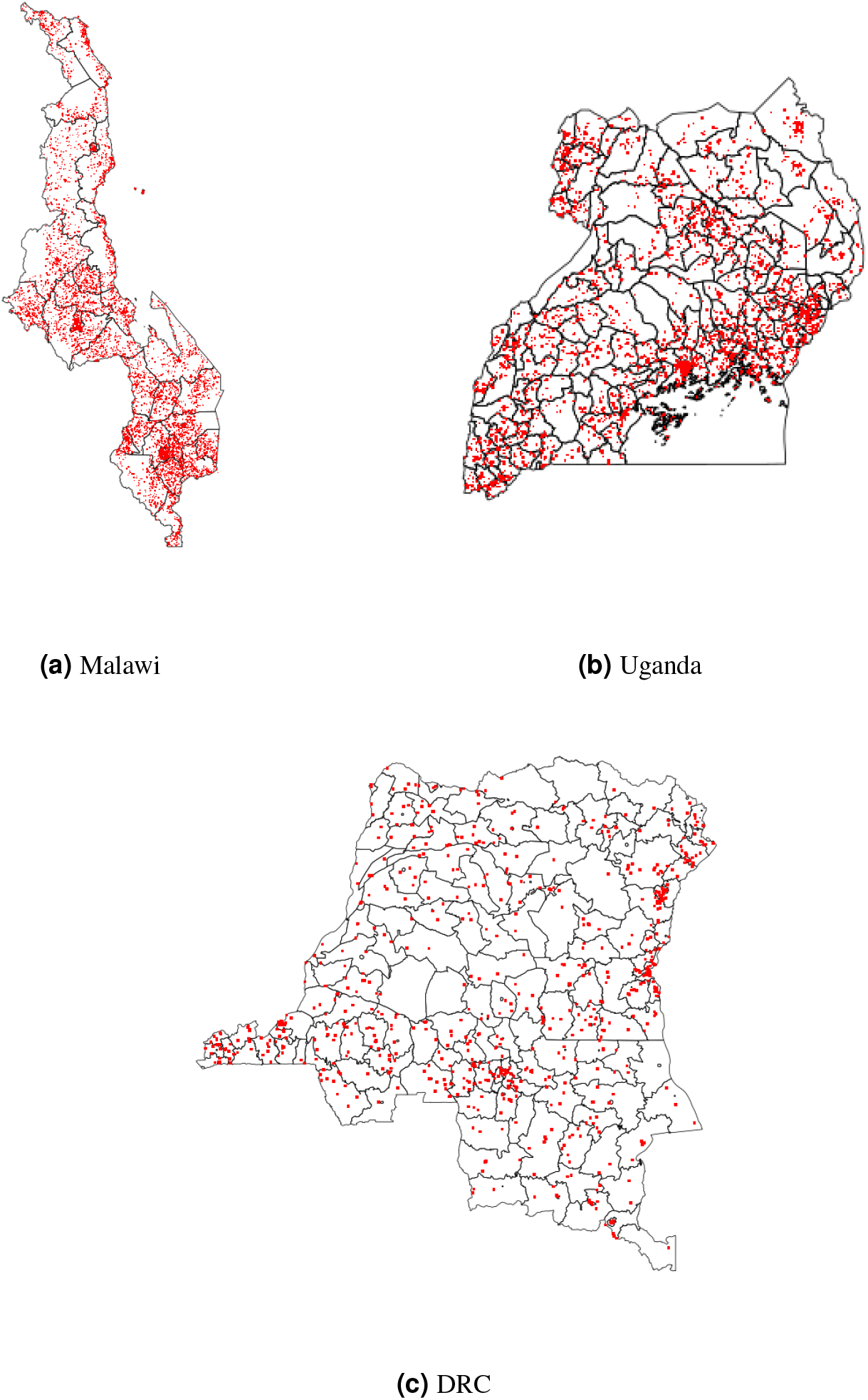
Spatial distribution of survey clusters (observed data) across all years. Size of the red dots does not scale with data richness

**Figure 2.**
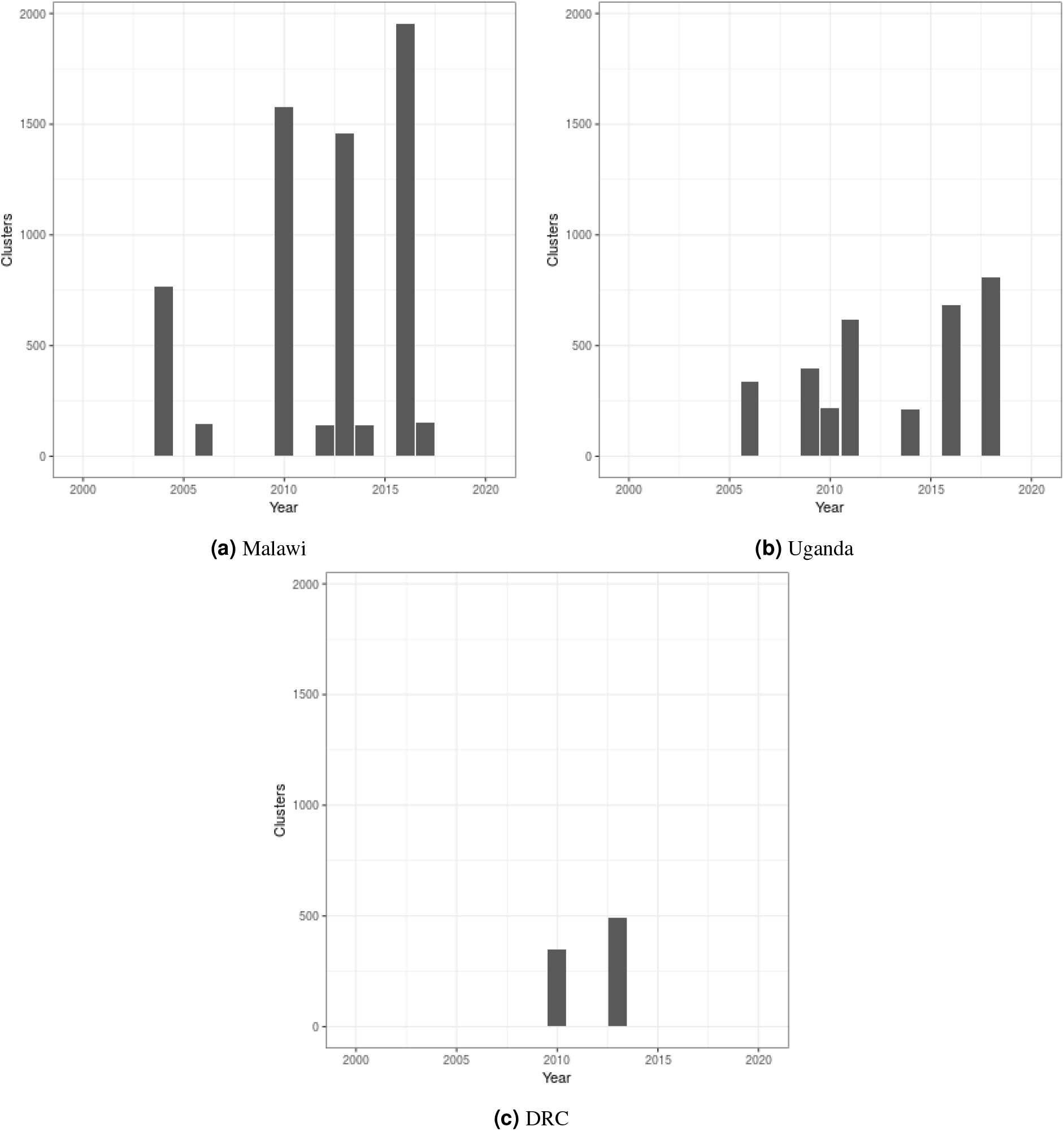
Number of survey clusters (enumeration areas) available for cattle density mapping in each year and country. Results were nearly equivalent for pigs

### Model selection (SPDE only)

Leave-one-out cross-validation (LOOCV) results are presented in Figure 3; we have removed models and surveys with very high mean squared error (MSE) for the sake of interpretability of these figures—as detailed in the accompanying caption—however these surveys and models were included when examining overall model performance. Model 6 performed best for Uganda (MSE 0.076), model 3 for Malawi (0.023), and model 2 for DRC (MSE 0.048).

**Figure 3.**
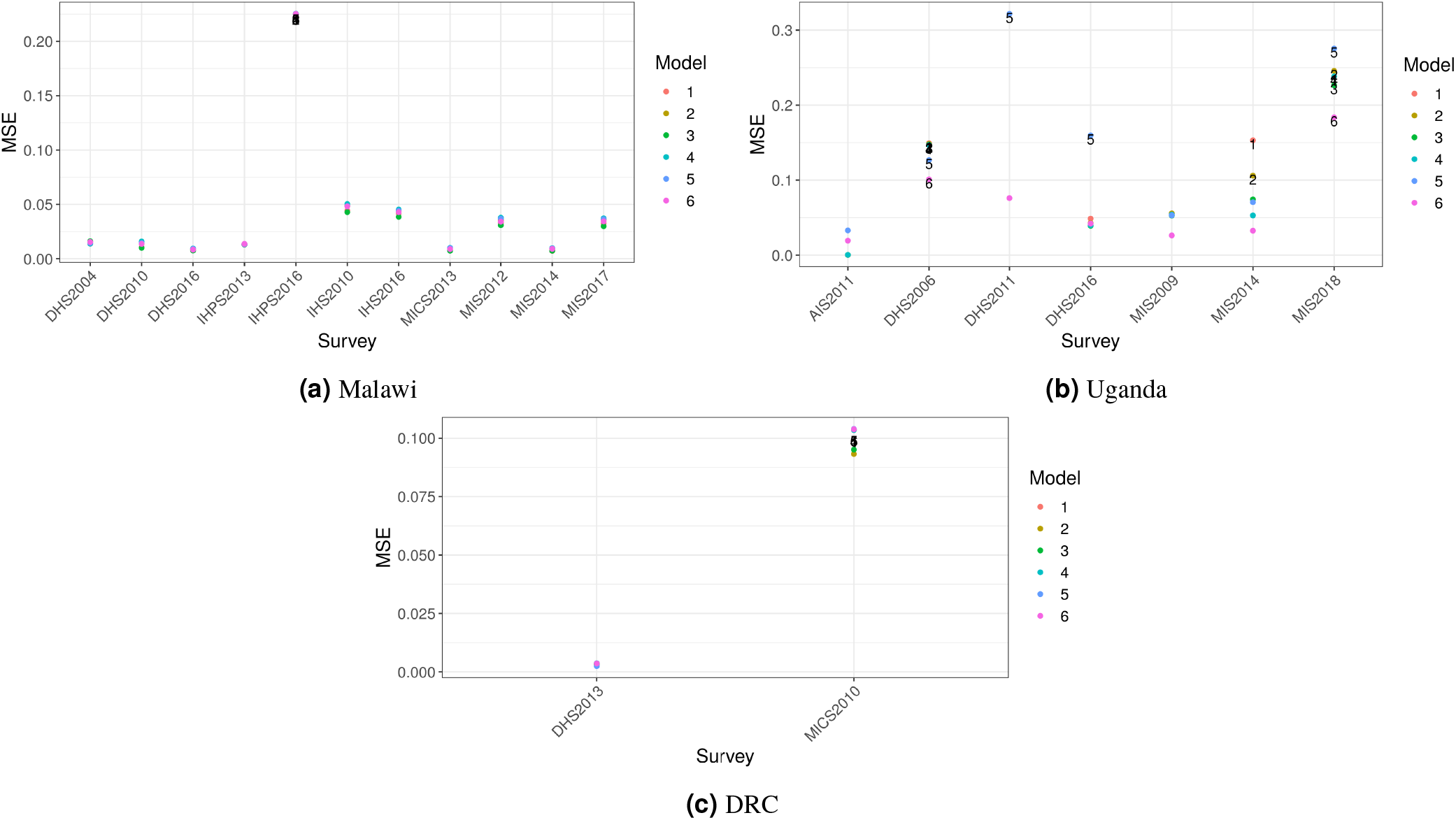
Leave one out cross validation results. Models with mean squared error (MSE) >0.1 are labeled. Removed from figures (but not MSE calculations) for interpretablity: (a) Malawi: 2004 Integrated Household Survey, all models; Uganda (b): 2009 National Panel Survey (NPS), all models, 2011 DHS, models 1-2, 2010 NPS, models 1 and 3-6, and 2011 NPS, models 1-3 and 5

In all three countries, we found extremely strong positive associations between our livestock density estimates and those derived from GLW-3 (coefficients greater than 100), for both cattle and pig density. These remarkably high coefficients can be explained by the coarser resolution of GLW-3 compared with our maps and WorldPop data (0.083° versus 0.017° and 0.0083° pixels, respectively), resulting in markedly higher density estimates for GLW-3 when we calculated density on these data as detailed in the Methods section. They do, however, indicate general agreement between our estimates and those of GLW-3.

### Maps

Density maps for 2005 and 2010 for Malawi and Uganda, 2010 and 2015 for DRC, and for 2008 for South Sudan, are presented in Figures 4-10. Results (median and width of the posterior 95% credible interval) as .tif files for every year for Malawi, Uganda, and DRC, and as shapefiles for 2008 for South Sudan, are available in a GitHub repository: https://github.com/JulianneMeisnerUW/LivestockMaps. Providing estimates of both median and uncertainty allows users to assess the quality of the estimates for each species, country, and year.

In Malawi (Figures 4-5), cattle density was highest in the northern and southern extents of the country, and pig density generally decreased along a west-east gradient. Across all years and clusters, median cattle density was 0.05, and median pig density was 0.07. Per FAOSTAT total stock estimates^13^ and World Bank national population estimates,^14^ mean national cattle density over the period 2000-2019 was 0.08, and mean national pig density was 0.15.

**Figure 4.**
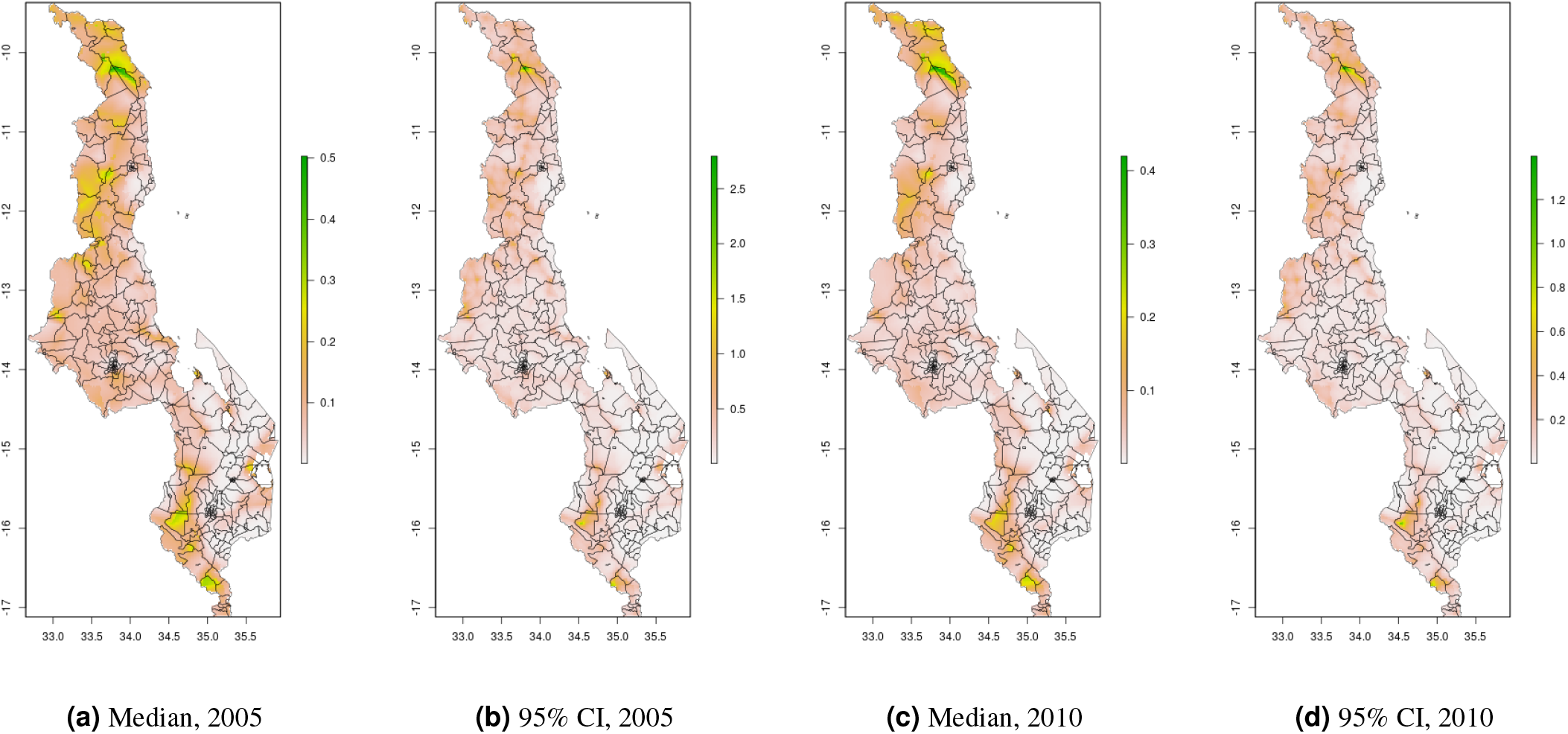
Cattle density mapping results, Malawi, 2005 and 2010. Median (a, c) and width of posterior 95% credible interval (b, d)

**Figure 5.**
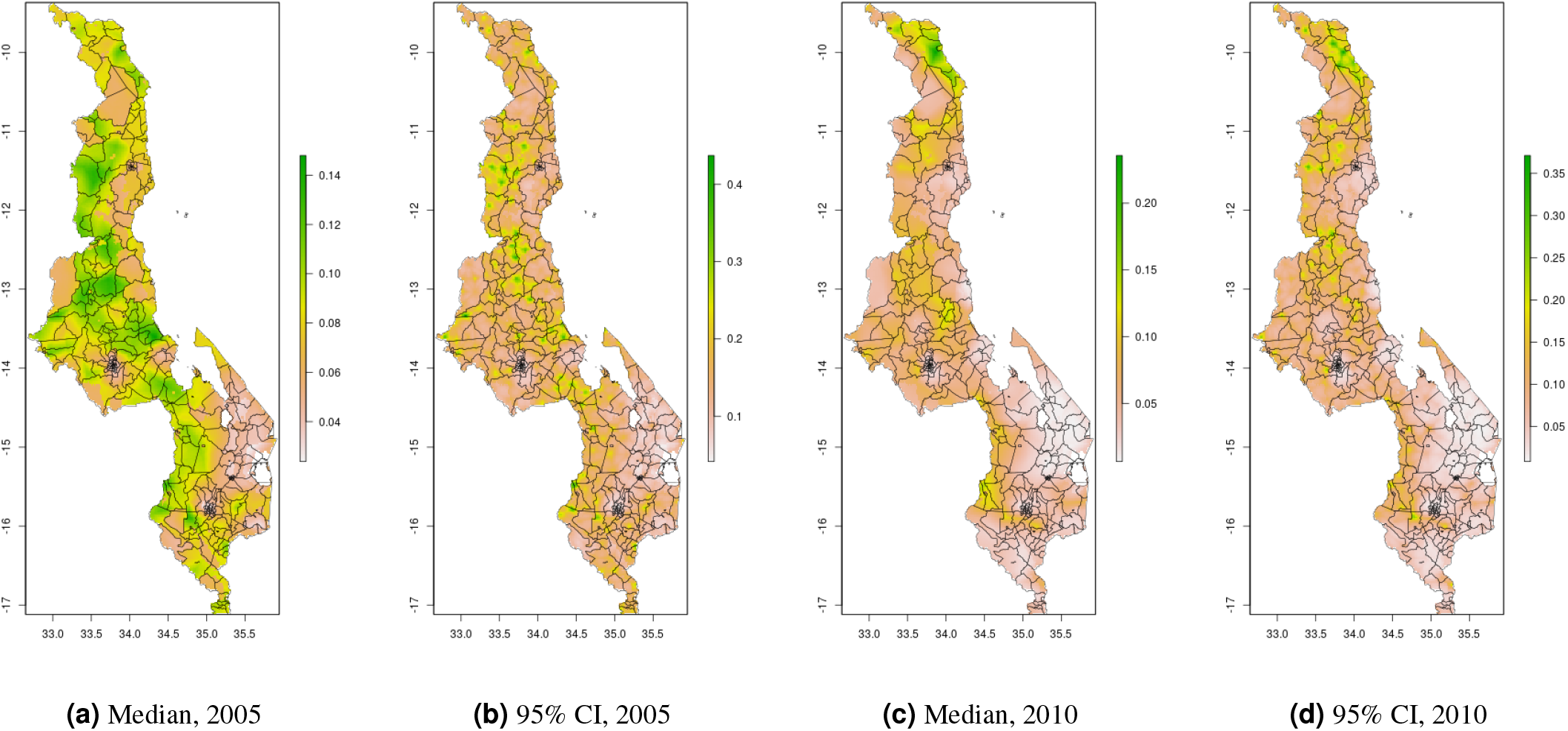
Pig density mapping results, Malawi, 2005 and 2010. Median (a, c) and width of posterior 95% credible interval (b, d)

In Uganda (Figures 6-7), the “cattle corridor” is clearly visible in both years, while pig density appears to be higher in the south of Uganda. Across all years and clusters, median cattle density was 0.23, and median pig density was 0.02. Per FAOSTAT total stock estimates^13^ and World Bank national population estimates,^14^ mean national cattle density over the period 2000-2019 was 0.33, and mean national pig density was 0.07.

**Figure 6.**
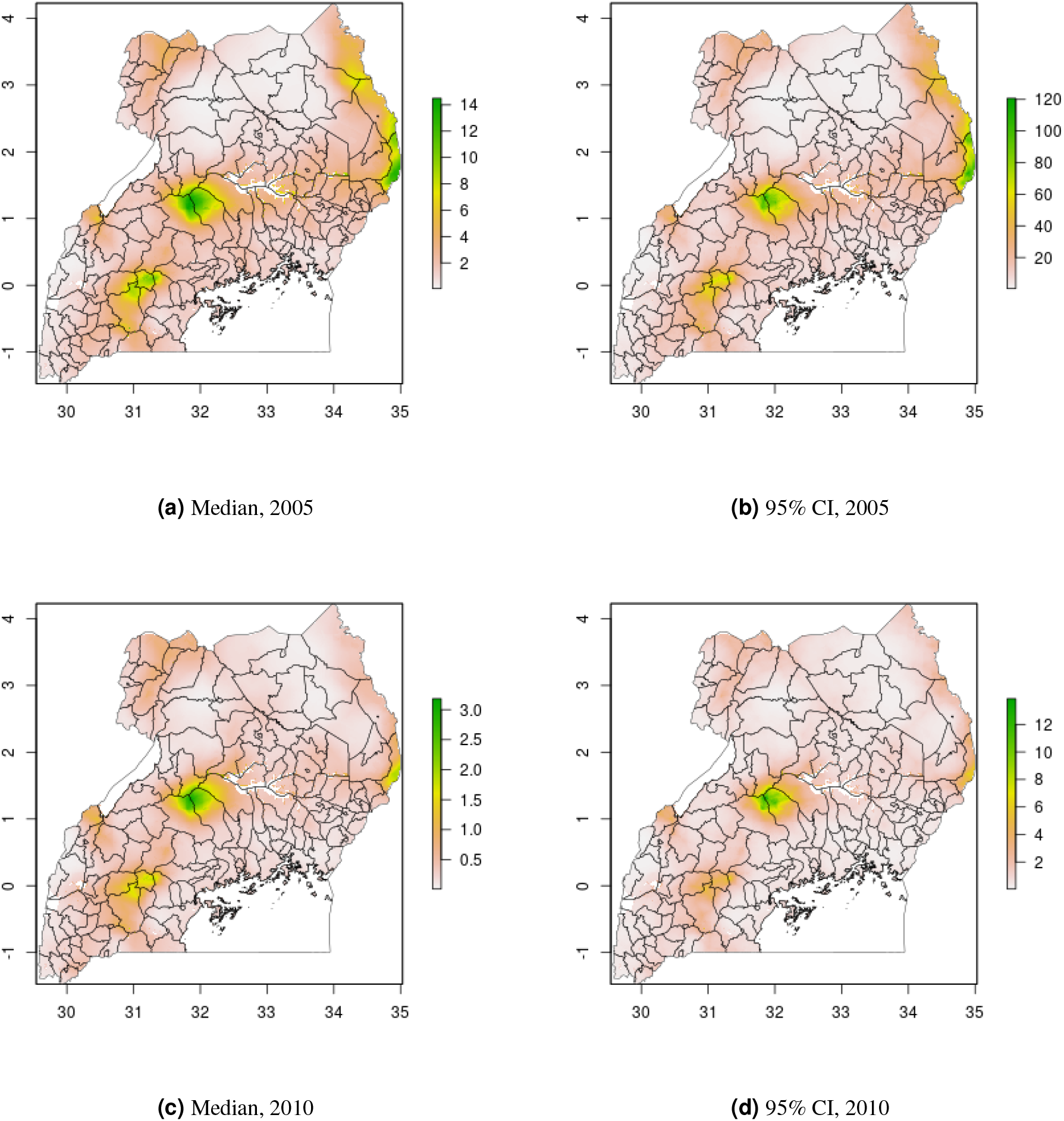
Cattle density mapping results, Uganda, 2005 and 2010. Median (a, c) and width of posterior 95% credible interval (b, d)

**Figure 7.**
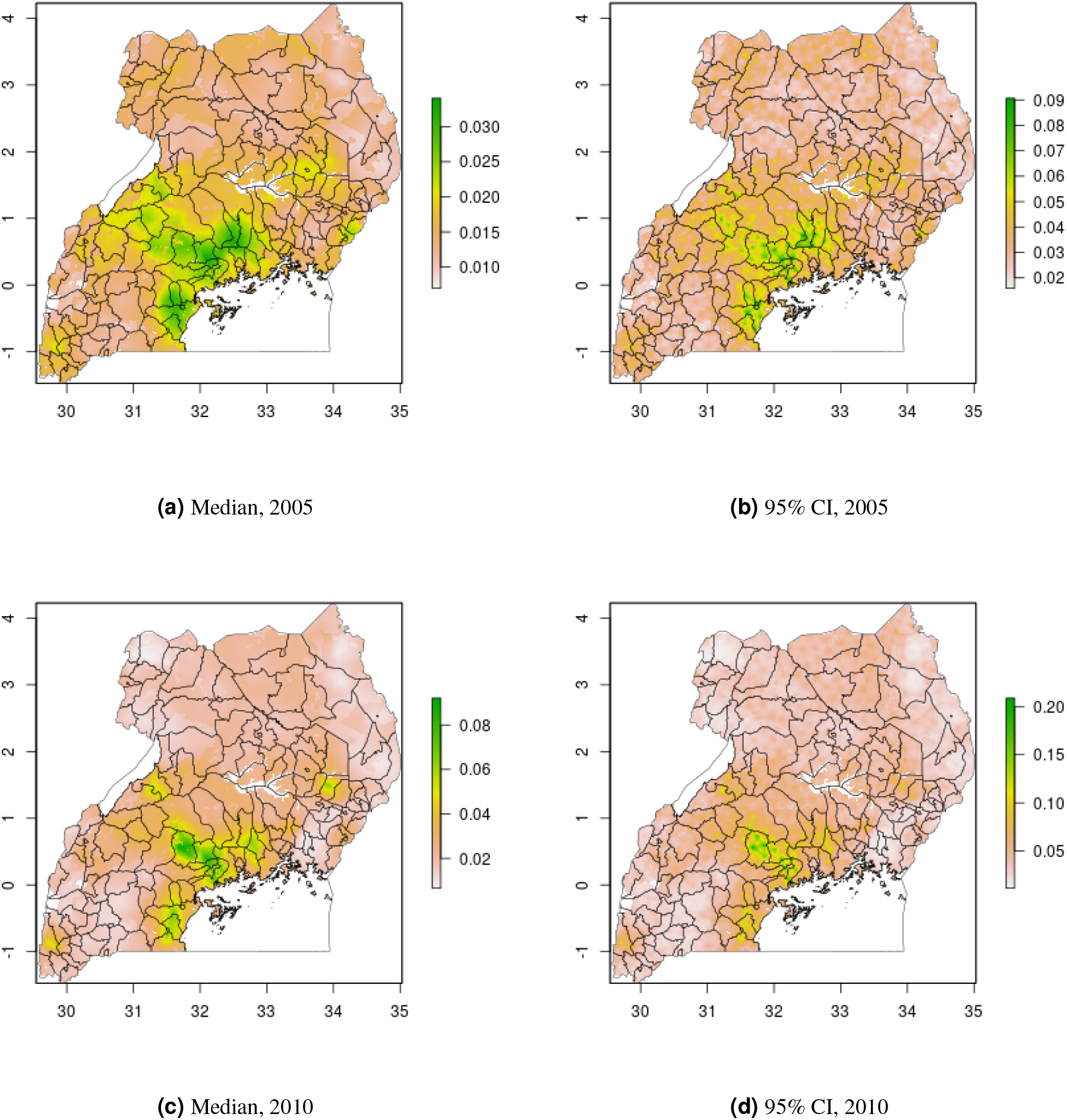
Pig density mapping results, Uganda, 2005 and 2010. Median (a, c) and width of posterior 95% credible interval (b, d)

In DRC, there are “hotspots” of cattle density in the east and southwest of the country, and no obvious spatial patterns for pig density (Figures 8-9) Per FAOSTAT total stock estimates^13^ and World Bank national population estimates,^14^ mean national cattle density over the period 2008-2015 was 0.013 for cattle and 0.014 for pigs, while across all years and clusters our median estimate was 0.037 for cattle and 0.053 for pigs.

**Figure 8.**
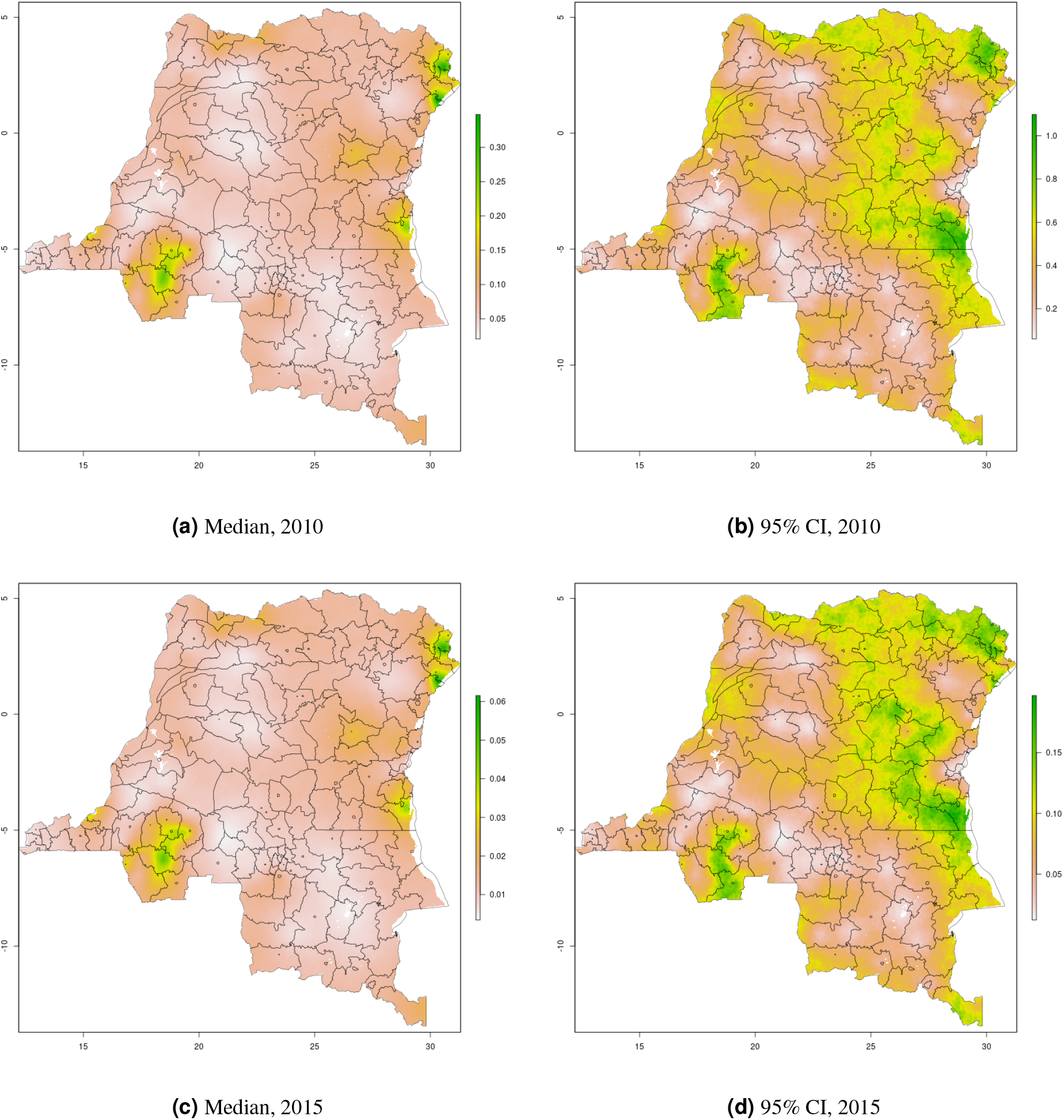
Cattle density mapping results, DRC, 2010 and 2015. Median (a, c) and width of posterior 95% credible interval (b, d)

**Figure 9.**
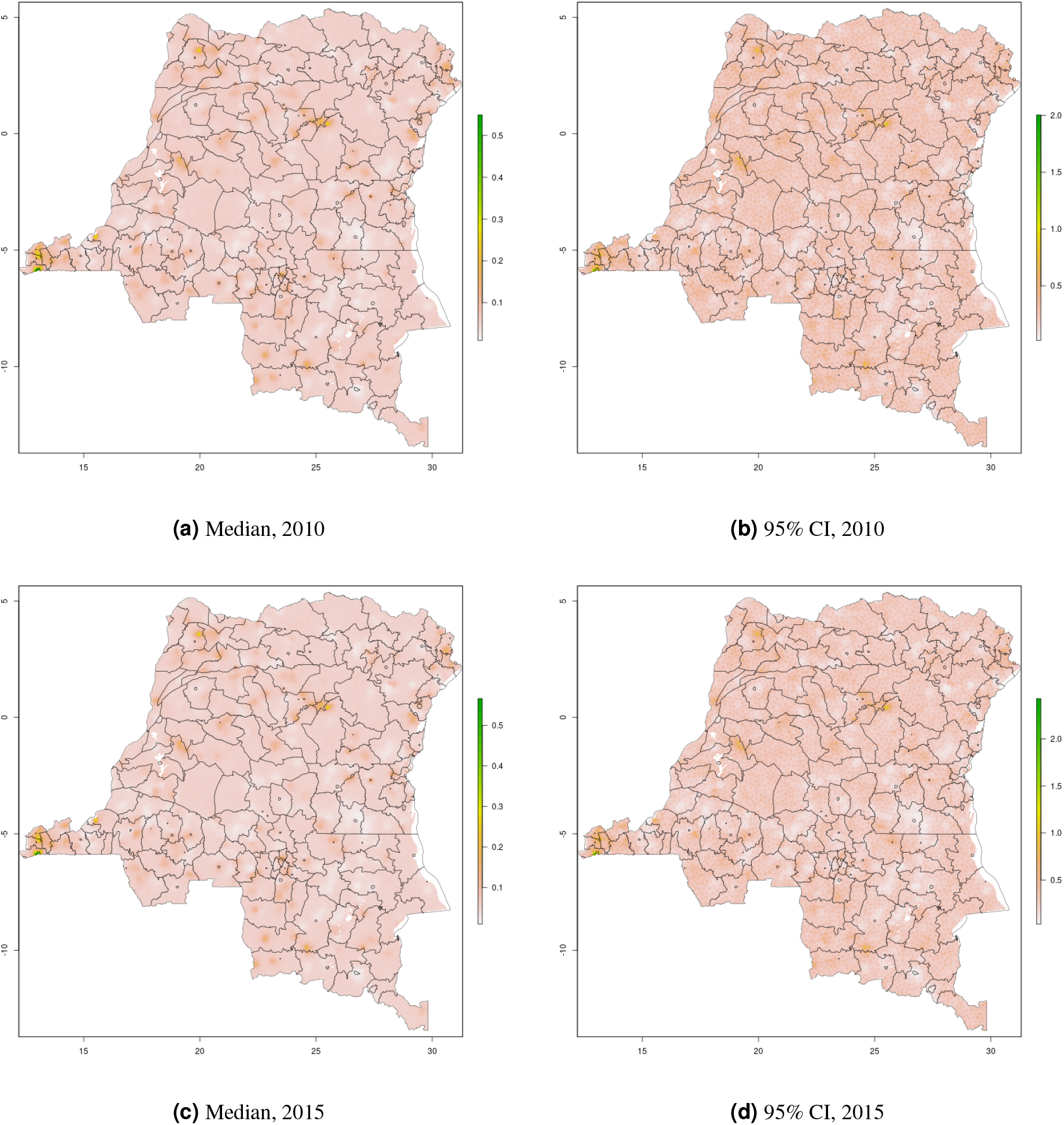
Pig density mapping results, DRC, 2010 and 2015. Median (a, c) and width of posterior 95% credible interval (b, d)

In South Sudan (Figure 10), cattle density was on average higher than pig density, and there is a clear decreasing southeast-northwest trend in cattle density. For pig density in South Sudan, other than generally low densities in the northwest of the country, there are no clear spatial trends. Across all counties, median density was 1.56 for cattle and 0.007 for pigs in 2008. Per FAOSTAT total stock estimates^13^ and World Bank national population estimates^14^, mean national cattle density over the period 2000-2019 was 1.33; no FAOSTAT entry is available for pigs in South Sudan.

**Figure 10.**
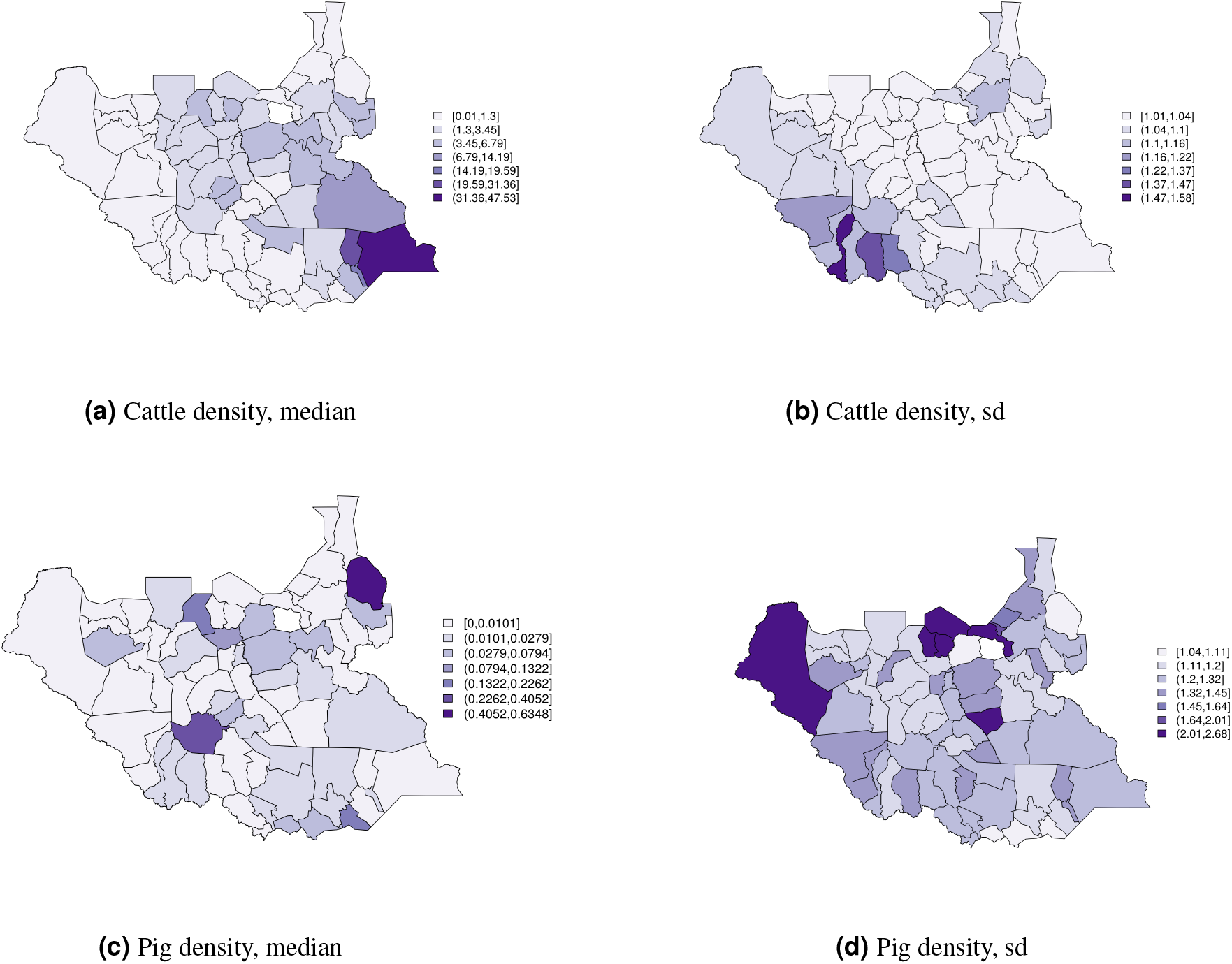
Cattle (a-b) and pig (c-d) density mapping results, South Sudan, 2008. Median (a, c) and standard error (b,d)

## Discussion

We have generated a set of cattle and pig density maps for Malawi, DRC, Uganda, and South Sudan, publicly available for use in economic, public health, environmental, agricultural, and other applications. Our results compliment GLW-3 efforts by producing a longitudinal product in Malawi, DRC, and Uganda, parameterizing livestock as the ratio of heads of cattle and pigs to the human population, and utilizing distinct input data and an entirely different modeling approach.

In contrast with GLW-3, we did not mask any pixels as unsuitable for livestock. The result is our findings impose no assumptions with regards to livestock presence in urban and high-altitude pixels, livestock incursion into protected areas, and wet versus dry season changes in the distribution of water bodies, but may result in bias in pixels corresponding to large and permanent bodies of water.

There are several limitations to our results. First, there may be inaccuracies in the input data, for instance over-reporting of livestock ownership due to the social desirability of owning a larger herd, or under-reporting due to concerns about increased taxation. Other inaccuracies may arise due to jittering in the input data, in which clusters in publicly-available datasets are randomly displaced to protect participant privacy.^15^ While ground-truthing is both cost-prohibitive and impractical for a longitudinal product, our LOOCV results indicate low MSEs overall, and good agreement between our results and GLW-3 results is a promising indicator of the validity of both products. Furthermore, our findings generally agreed with national density estimates, and we would not expect perfect agreement with such estimates as local distribution patterns for humans is unlikely to be equivalent to local distribution patterns for livestock and we are comparing medians to means. In addition to jittering in the input data sources, misclassification of livestock density may arise from the location definition used in our models, whereby livestock are assigned to the centroid of the corresponding enumeration area. Thus our maps should be interpreted as the density of livestock to humans when both are in their place of residence (i.e., at nighttime). While this may be a poor reflection of livestock presence in pastoralist systems, it is difficult to conceptualize any other definition of livestock density as the daily movements of humans and livestock may be distinct.

Second, our results treat livestock production systems as equivalent within a given species, however the environmental, socioeconomic, and public health relevance of production system types (e.g., pastoralist vs. intensive) are distinct, potentially reducing the utility of our results for select uses. Furthermore, users will need to take care to ensure the predictors (spatial covariates) used in our SPDE models do not introduce circularity in their analyses.

Third, our approach to generating urbanicity surfaces reflects the challenging nature of meaningfully assigning unsampled cluster-years as urban versus rural. Our approach performed well in the test data with the exception of urban clusters in Uganda, for which it performed very poorly. Alternative approaches, such as using Global Human Settlement Layer data^16^ and other existing urbanicity surfaces, do not necessarily correspond to the definition of urbanicity needed in our application, nor are they generally available as a longitudinal product.

Fourth, due to limited data in DRC and South Sudan, we could not include a space-time interaction term in DRC—imposing the assumption that spatial trends are constant in time, and temporal trends are constant in space—and could only produce a county-level map for South Sudan, and only for a single year.

Finally, our estimates are accompanied by uncertainty which varies across time, countries, and clusters. Uncertainty was highest for DRC and for early years, rendering our maps less useful in these cluster-years. It is our hope that by producing uncertainty estimates alongside of medians, it will be transparent to users where our results are more or less stable.

For poor livestock keepers throughout the world, livestock represent a lifeline, providing financial security, nutrition, transportation, and connection to cultural identities. Livestock, however, also represent a source of exposure to zoonoses, and a driver of environmental change on local and global scales. To grapple with these divergent effects in research and policy, high-resolution estimates of livestock distribution tethered to human population distribution are needed. We present such a product, which, despite its limitations, represents an important extension to GLW-3 with high demonstrated validity and broad potential utility.

## Methods

### Data collection and processing

Data sources were identified by searching GHDx and the IHSN Central Data Catalog,^17, 18^ and microdata were downloaded from publicly-accessible websites or by request to relevant national agencies. Both Demographic and Health Survey (DHS) data and population and housing census data were downloaded from IPUMS;^19, 20^ most other surveys were downloaded from links in the relevant IHSN entry. Data were processed by source; data sources by country are detailed in Table 1.

**Table 1.**
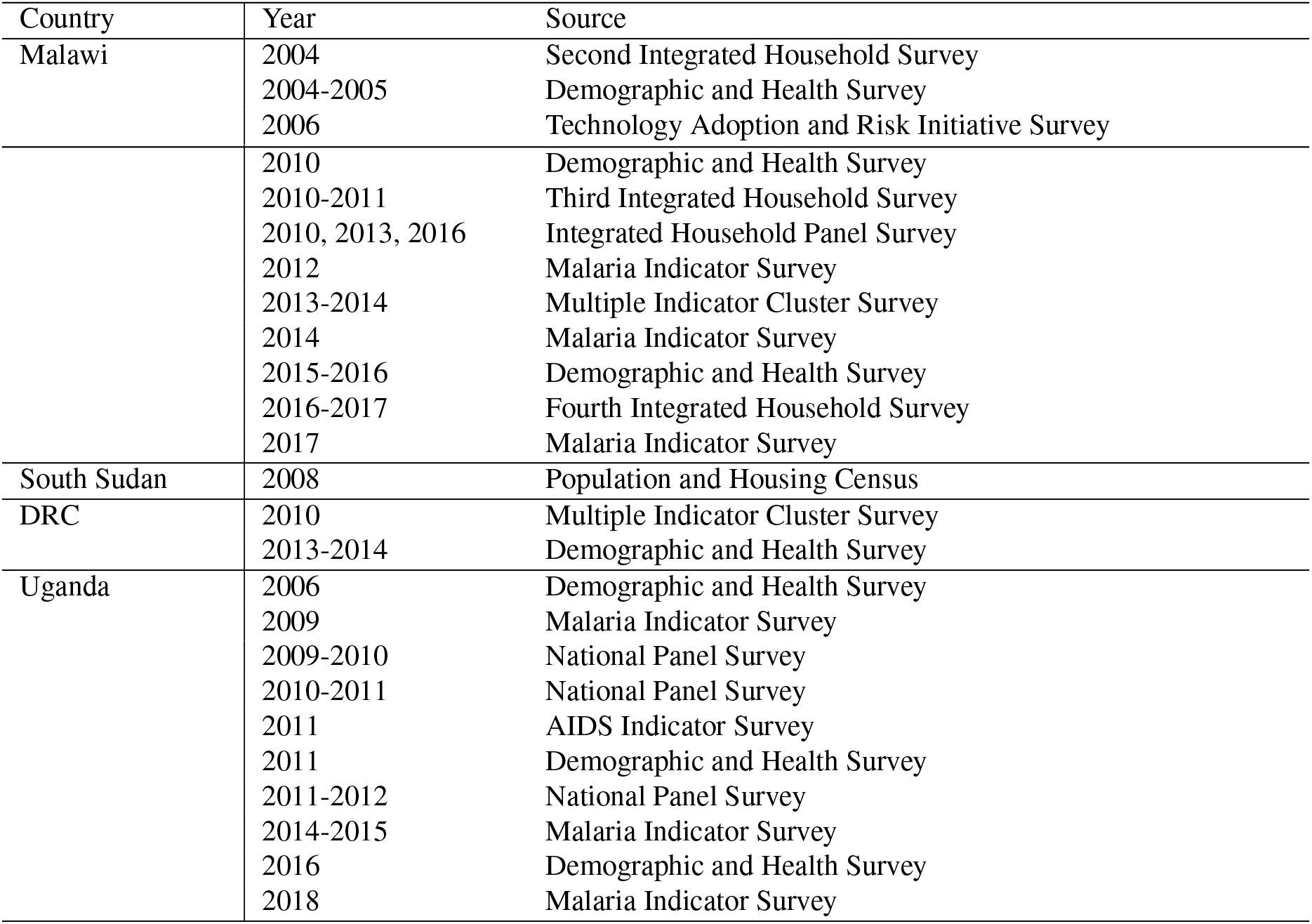
Data sources by country

As data were only available for a single year for South Sudan (2008), no longitudinal mapping could be performed for this country. Furthermore, these data provided geolocation only to the level of the county (administrative level 2), thus we have used small area estimation, rather than the SPDE approach, to generate county-level maps in South Sudan. In Malawi, Uganda, and DRC, survey cluster—the smallest geographical sampling unit of a household survey, typically an enumeration area consisting of approximately 100 households—was the unit of analysis.

### Model fitting

#### Malawi, Uganda, and DRC

For Malawi, Uganda, and DRC, we used the SPDE approach to Gaussian process (GP) modeling. The goal of this approach is to smooth over spatial point data and generate a complete surface.

##### SPDE approach

A spatial process *S*(s) is a GP if the joint distribution of *S*(s_1_),…, *S*(s_*n*_) (over the whole study area) is an *n* dimensional Gaussian distribution for any integer *n* and any set of locations s_*i*_ (e.g., defined by latitude and longitude). That is, the spatial process at each location has a Gaussian distribution.

In Bayesian hierarchical spatial modeling, spatial dependence is generally represented by a precision matrix (inverse of a variance-covariance matrix), with the Matérn family being a popular choice of covariance function.^21^ It can be shown that Matérn GPs can be represented as the solution an SPDE, but for a GP with *n* clusters, a *n* dimensional normal distribution must be modeled, which quickly becomes computationally intractable. However, under certain conditions the resulting Gaussian random field has the continuous Markov property; i.e., it is a GMRF (Gaussian Markov random field). This means a particular location’s spatial process depends only on that of its neighbors, not the entire map, resulting in a “sparse” precision matrix (many 0 entries) and markedly simplifying computation.^22^ In our application of the SPDE approach, we parameterize the Matérn in terms of three parameters: 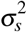 (spatial variance), *ρ_s_* (spatial range), and *v* (shape, or smoothness).

Now that we have represented the GP using an SPDE, the next step is to approximate continuous space using a weighted combination of basis functions, where the weights are picked to have a sparse precision matrix. In `R-INLA` this is done using a set of non-overlapping triangles which together comprise a mesh over the study region. The basis functions are piecewise linear, equal to 1 on a given vertex and 0 on all other vertices. If one imagines elevating a single mesh vertex, a three-sided pyramid is formed, and the discrete approximation to the GP is a weighted combination of these pyramids. A projection matrix *A* projects from the mesh verticles to the *n* study locations (here, clusters).

Choice of the mesh dictates the resolution of the spatial effect, with a rule of thumb that features more than two triangles large are resolved well, while features smaller than a triangle will be biased in proportion with the triangle size.^21^ Our meshes for Malawi, Uganda, and DRC are presented in Figure 11 below. In all three countries, we set our mesh to have an inner and outer mesh, moving boundary effects away from the study area (i.e., country borders). We set the triangles to be smaller in the inner mesh (maximum edge length of 0.3°) than the outer mesh (maximum edge length of 0.6°), and the extension to be 1°.

**Figure 11.**
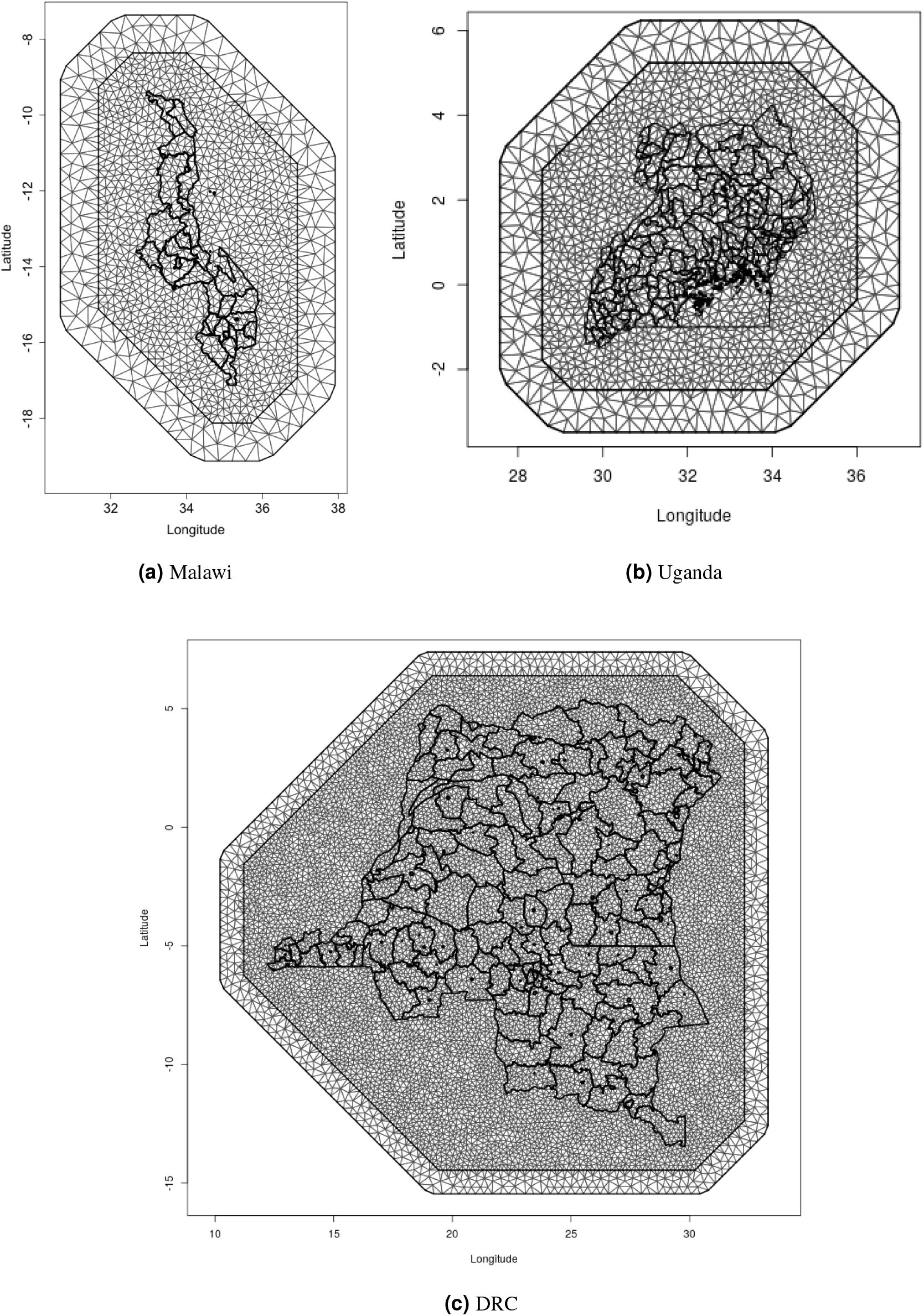
Meshes used for SPDE

##### Predictors

When the data are complex survey data, the sampling scheme may be accounted for by including design variables— district and urban/rural status in this case—in the regression model. As district is confounded by space, the spatial random effect (detailed above) accounts for this aspect of study design, and district was not included as a predictor in our models.

Predictors included urban/rural status (binary), protected areas (binary), elevation, and bodies of water (binary). While there are many definitions of urbanicity, as urban/rural status is a design variable, the desired definition for this predictor is the value that a given cluster would take had it been sampled by a household survey in a given year. As the data sources we used for livestock mapping largely used the most recent population and housing census as their sampling frame, for each country we generated a 1km^2^ urban/rural surface for each sampling frame year. We accomplished this using logistic regression models, with population density from WorldPop^23^ and nighttime lights from the National Oceanographic and Atmospheric Administration’s Nighttime Lights Time Series,^24^ as the predictors. For validation, we split the data into a 2/3 training and 1/3 test set, and evaluated model performance on the test set. We found these models performed very well for rural clusters (95% correct for Uganda, 96% for Malawi and DRC), however performance varied across countries for urban clusters (43% for Uganda, 86% for Malawi, 78% for DRC). As the vast majority of clusters in all three countries are rural, overall performance remained adequate.

Data on protected areas came from the World Database on Protected Areas;^25, 26^ while this database is updated monthly, there is no publicly-accessible archive. For bodies of water we used levels 1 (lakes ≥ 50km and reservoirs ≥0.5km) and 2 (permanent open water bodies with a surface area of ≥0.1km) data from the Global Lakes and Wetlands Database.^27^ Finally, we used 7.5-arc-second data—which have a root mean squared error of 26-30 meters—and median elevation from GMTED2010 28 10 for elevation.^28^ Note, in contrast with GLW-3 these predictors were used to improve model fit, not to mask unsuitable pixels.^10^

##### Models

We fit a three stage Bayesian hierarchical model as follows for the minimal model, fit separately for cattle and pigs, and separately for each country:

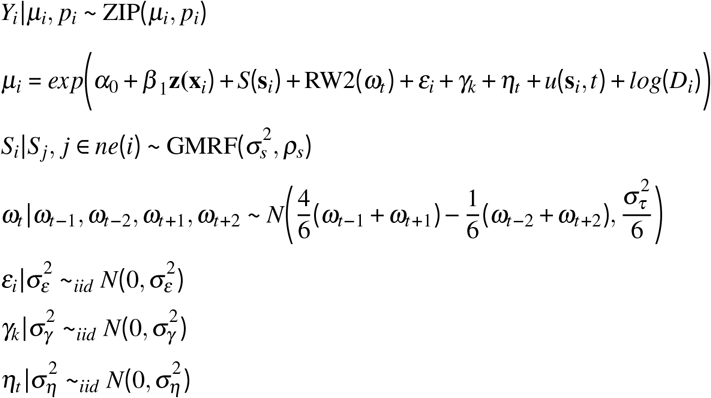

Where:

- *i* indexes cluster
- *p_i_* is the hyperparameter that pertains to the model for the zeroes for cluster *i*
- *Y_i_* is the number of livestock in cluster *i*
- *D_i_* is the number of humans in cluster *i* (offset)
- *α*_0_ is the intercept
- *β* is a vector of coefficients for the predictors
- **z**(**x**_i_) is a vector of predictor variables for location *i*
- *S*(**s**_*i*_) are the spatial random effects (error terms), assumed to follow a Markovian Gaussian random field (GMRF) with variance parameter 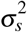 and range parameter *ρ_s_*, and *ne*(*i*) are the neighbors of cluster *i*.
- *ω_t_* is a random walk 2 (RW2) random effect on time (years)
- *ε_i_* is an unstructured cluster-level random effect (nugget)
- *γ_k_* is an unstructured random effect on survey
- *η_t_* is an unstructured random effect on time (years)
- *u*(**s**_*i*_, *t*) is a space-time interaction which is approximated by an SPDE in space combined with an AR(1) process in time

We fit these models as type 1 zero-inflated Poisson (ZIP) models in `R-INLA`,^29^ which combine a distribution for the proportion of zeros with the Poisson distribution. The type 1 likelihood is given as:

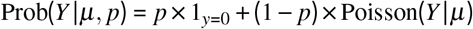

where the first part is the process that generates the zeros (i.e., clusters with no livestock; 1_y_i_=0_ is an indicator that cluster *i* has 0 livestock), and the second part generates the livestock counts in cluster *i*. This likelihood is a mixture of structural zeros (*p_i_*) and sample zeros (1 - *p_i_*).^30^

For model selection, we fit a series of five models with equivalent random effects but varying fixed effects: intercept only (model 1); intercept and urban/rural (model 2); intercept urban/rural, and protected area (model 3); intercept, urban/rural, protected area, and bodies of water (model 4); and intercept, urban/rural, protected area, bodies of water, and elevation (model 5). We also fit a sixth model with an RW1 effect in time, rather than RW2, and all fixed effects included in model 5.

In DRC, due to the limited data availability, we fit models only from 2008-2015 (i.e., extrapolating 2 years in each direction from the input data), and we restricted random effects to the structured (SPDE) effect on space, the structured (RW1 or RW2, depending on the model) effect on time, and the unstructured (iid) effect on space (cluster). That is, the space-time interaction and unstructured random effects for survey and time were removed.

##### Priors

As in Wakefield et al.,^31^ for the spatial random effect we assigned a fixed shape *v* = 1, and a “penalized complexity” (PC) prior^32, 33^ for the spatial range *ρ_s_* and marginal standard deviation *σ_s_*, such that *Pr*(*ρ_s_* < 0.3) = 0.05, and *Pr*(*σ_s_* > 1) = 0.05. For the spatial range parameter, this prior can be interpreted as the 5% quantile corresponds to 0.3°, which is approximately 4% and 9% of the extent of Malawi in the north-south and east-west directions, respectively, and approximately 5% and 1.5% of the extent of Uganda and DRC in each direction, respectively. This spatial range, as well as the mesh, were chosen to be adequately fine to allow for construction of a high-resolution raster map, but adequately coarse to account for jittering of cluster locations in the source data (a means to protect privacy) and to make the model computationally feasible. For the marginal standard deviation this prior corresponds to a posterior 95% credible interval for the residual rate ratio—deviations in livestock density above or below the mean model, attributable to the random effect in question—of (0.37, 2.72). We used `R-INLA`’s default priors for the structured temporal effects (RW1 and RW2).

For the AR(1) process in time, again using the PC prior specification we set *Pr*(*ρ_t_* > 0.5) = 0.8 as the prior. Finally, for the precision (inverse of variance) parameters for each of the independent and identically distributed (iid) random effects 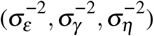, we specified the prior as *Pr*(*σ*^2^ > 0.5) = 0.01, which results in a posterior 99% credible interval for the residual variance of (0.6, 1.6).

##### Prediction on a regular grid

After fitting our models, we projected livestock density on regular grid, generating 0.017° x 0.017° grid cells in each country (Figure 12). This choice governed the resolution of our final raster maps.

**Figure 12.**
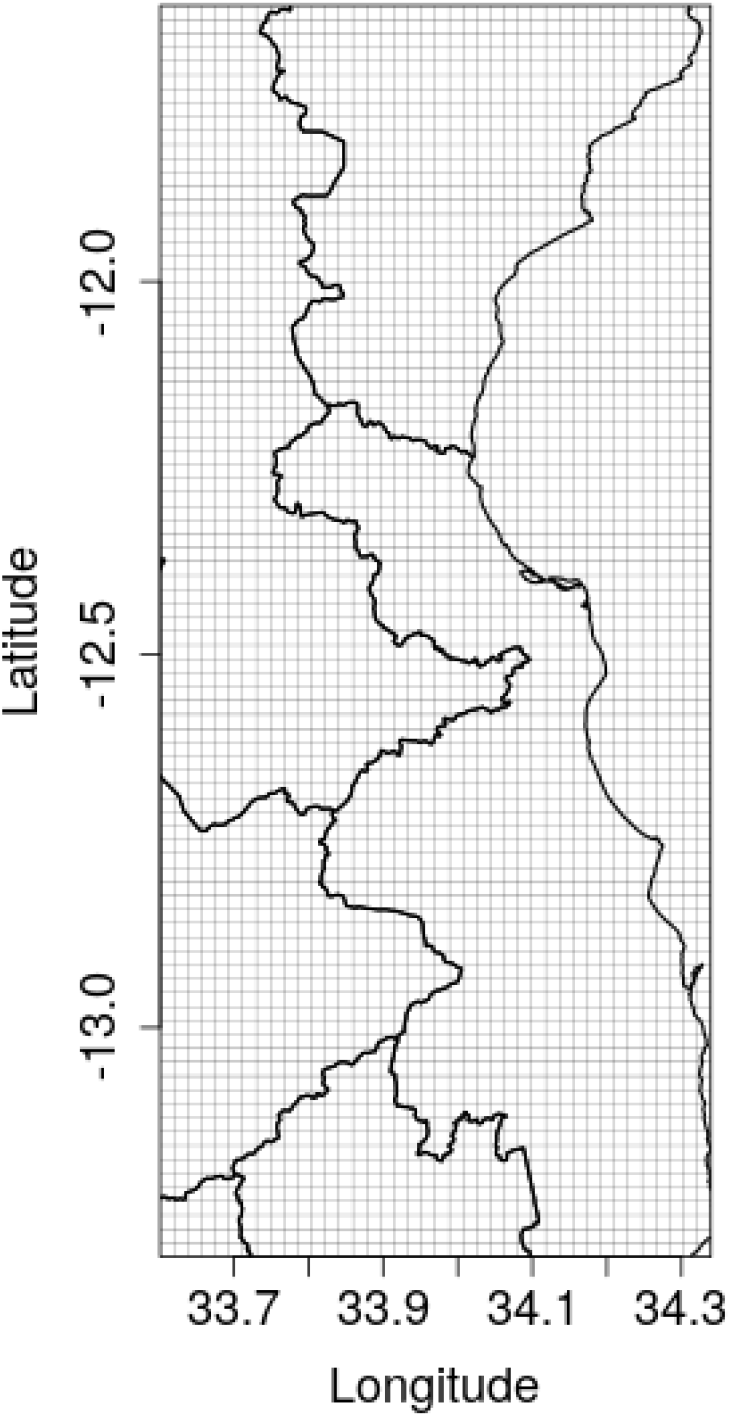
Regular grid for prediction, cropped to the northwest of Malawi. Prediction grids were of equivalent resolution in Uganda and DRC

We did not use any of the iid (unstructured) random effects for prediction as these were assumed to reflect measurement error. Thus, prediction was comprised of each model’s linear predictor, a vector of predictor values for each grid cell, the structured spatiotemporal random effects, and an appropriate projection matrix for each random effect. To produce estimates of uncertainty, we took 1,000 draws from the posterior distribution of the structured random effects using the command `inla.posterior.sample()`, and present here the median and width of the posterior 95% credible interval over these 1,000 estimates.

Because we did not use an offset in our predictions, we essentially “flattened” the distribution of the human population over the map: our final estimate is that of livestock counts in a grid cell with 1 human being, equivalent to the ratio of livestock to humans, which we are calling livestock “density.”

##### Model selection

After fitting our models, we performed model selection via LOOCV for the cattle maps by survey, as we could most readily conceptualize the missing data as hypothetical surveys which were not conducted or samples not collected by a given survey. Each fold left out a random 25% sample, distributed evenly among sampling strata defined by district and urban-rural status. We present model performance as MSE, which we calculated as:

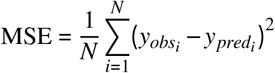

where *y_obs_i__* is the observed density at location *i*, and *y_pred_i__* is the predicted density at location *i*.

##### External validation

For external validation we regressed livestock density estimated from GLW-3 (dasymetric product), using WorldPop data as the denominator, on our final 2010 estimates of livestock density. ^23, 34, 35^

We also compared median density estimates, taken over all years for each country, to median estimates derived from national-level FAOSTAT total stock estimates^13^ and World Bank national (human) population estimates,.^14^

#### South Sudan

##### Weighted estimates

After reading in the data, we used the `svyby()` and `svyratio()` functions in the `survey` package in R^36^ to generate weighted estimates of county-level density, using sample weights contained in the IPUMS subset^20^ and the ratio estimator below, where *i* indexes household and *c* indexes county:

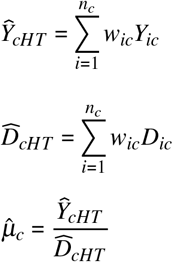

wheres

- *Y_ic_* is the number of livestock in household *i*, county *c*, and 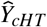 is the Horwitz-Thompson estimate for the number of livestock in county *c*
- *D_ic_* is the number of residents in household *i*, county *c*, and 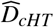 is the Horwitz-Thompson estimate for the (human) population in county *c*
- *n_c_* is the total number of households in county *c*
- *w_ic_* is the sampling weight for household *i* in county *c*, which is the inverse of sampling probability

We also used the `survey` package to estimate design-based standard errors for livestock density. Adapted from Mercer et al. 2014,^37^ we then specified 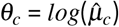, which, by the delta method, gives us the asymptotic sampling distribution:

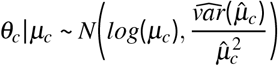

##### Smoothing

We performed spatial smoothing to stabilize the variance of the direct estimates by fitting the second and third stages of the three stage Bayesian hierarchical model as:

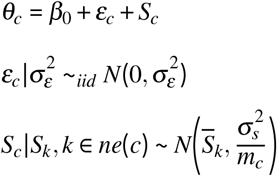

where

- *ε_c_* are county-level iid (unstructured) random effects with variance 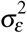
- *S_c_* are county-level structured random effects which follow the ICAR model with marginal variance 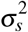
- *ne*(*c*) denotes neighbors (shared boundary) of county *c*
- *m_c_* is the number of neighbors of county *c*

and ICAR is the intrinsic conditional autoregressive model, which smooths each county’s random effect to that of its neighbors, with more smoothing performed for counties with fewer neighbors.

##### Priors

As for the SPDE models, we used PC priors for the smoothing model, with *Pr*(*σ_ε_*) > 1 = *Pr*(*σ_s_*) > 1 = 0.01. This yields a posterior 95% credible interval for each random effect’s residual rate ratio of (0.36, 2.71).^32, 33^

##### Model selection

As there was only one candidate model for South Sudan, model selection was not performed.

## Data availability

While existing data use agreements do not allow direct sharing of input data, all data were downloaded from publicly-available sources, detailed in Table 1, with links included in our References section. All analyses were performed in R, and all code are available in the GitHub repository linked in the Results section.

## Ethics

As this work relied solely on pre-existing, de-identified data, it does not constitute human or animal subjects research. Per the University of Washington Human Subjects Division, review and approval by the University of Washington Institutional Review Board is not needed (IRB ID: STUDY00004648)

## Author contributions statement

J.M.: conceptualiztion, methodology, software, formal analysis, data curation, writing - original draft, writing - review & editing, visualization. A.K.: data curation, resources, writing - review & editing. M.L.: data curation, resources, writing - review & editing. E.M.M: data curation, resources, writing - review & editing. A.I.T.: data curation, resources, writing - review & editing. J.W.: methodology, software, writing - review editing, visualization. A.R.R: methodology, writing - review & editing. D.P.: methodology, software, writing - review & editing. J.M.: methodology, writing - review & editing. P.R: methodology, supervision, writing - review & editing, resources.

## Additional information

### Competing interests

The authors declare no competing interests.

## Notes

https://github.com/JulianneMeisnerUW/LivestockMaps

### Competing Interest Statement

The authors have declared no competing interest.

## References

1. Herrero, M., Havlik, P., McIntire, J., Palazzo, A. & Valin, H. African livestock futures: Realizing the potential of livestock for food security, poverty reduction and the environment in sub-Saharan Africa. Off. Special Represent. UN Secr. Gen. for Food Secur. Nutr. United Nations Syst. Influ. Coord. (UNSIC), Geneva, Switz. 188p (2014).

2. Grace, D. et al. Mapping of poverty and likely zoonoses hotspots. Int. Livest. Res. Inst. (2012).

3. Hermesh, B., Rosenthal, A. & Davidovitch, N. The cycle of distrust in health policy and behavior: Lessons learned from the Negev Bedouin. PloS One 15, e0237734, DOI: 10.1371/journal.pone.0237734 (2020).

4. Hermesh, B., Rosenthal, A. & Davidovitch, N. Boundaries and Politics. Monash Bioeth. Rev. 37, 22–37, DOI: 10.1007/s40592-018-0079-9 (2019).

5. Thornton, P. Livestock production: recent trends, future prospects. Phil. Trans. R. Soc. B 365, 2853–2867, DOI: 10.1098/rstb.2010.0134 (2010).

6. The Livestock Revolution. http://www.fao.org/WAIRDOCS/LEAD/X6115E/x6115e03.htm. Accessed 2 Nov 2018.

7. Steinfeld, H.., Gerber, P., Wassenaar, T. D.., Castel, V.. & de Haan, C. Livestock’s long shadow: environmental issues and options. FAO 5, 7, DOI: 10.1890/1540-9295(2007)5[4:D]2.0.CO;2 (2006).

8. Lunde, T. & Lintjorn, B. Cattle and climate in Africa: How climate variability has influenced national cattle holdings from 1961–2008. PeerJ (2013).

9. Wint, G. & Robinson, T. Grided livestock of the world 2007. Tech. Rep., FAO, Rome (2007).

10. Gilbert, M. et al. Global distribution data for cattle, buffaloes, horses, sheep, goats, pigs, chickens and ducks in 2010. Sci. Data 5, 1–11, DOI: 10.1038/sdata.2018.227 (2018).

11. Hankerson, B. et al. Modeling the spatial distribution of grazing intensity in Kazakhstan. PLoS ONE 14, DOI: 10.1371/journal.pone.0210051 (2019).

12. Jahel, C. et al. Mapping livestock movements in Sahelian Africa. Sci. Reports 10, DOI: 10.1038/s41598-020-65132-8 (2020).

13. FAOSTAT. Live Animals. urlhttp://www.fao.org/faostat/en/data/QA. Accessed 16 Feb 2021.

14. The World Bank. Population, total. https://data.worldbank.org/indicator/SP.POP.TOTL. Accessed 16 Feb 2021.

15. Demographic and Health Survey: Methodology - collecting geographic data. https://dhsprogram.com/What-We-Do/GPS-Data-Collection.cfm. Accessed 22 Oct 2018.

16. European Commission. Global Human Settlement Layer. https://ghsl.jrc.ec.europa.eu/index.php. Accessed 09 Apr 2018.

17. Global Health Data Exchange. Institute for Health Metrics and Evaluation. http://ghdx.healthdata.org/. Accessed 24 Aug 2017.

18. Central Data Catalog. Integrated Household Survey Network. https://catalog.ihsn.org/index.php/catalog. Accessed 24 Aug 2017.

19. Heger Boyle, M., King, M. & Sobek, M. IPUMS-Demographic and Health Surveys: version 7 [dataset]. Minnesota Population Center and ICF International, 2019, DOI:https://doi.org/10.18128/D080.V7.

20. Minnesota Population Center. Integrated Public Use Microdata Series, International: version 7.2 [dataset]. Minneapolis, MN: IPUMS, 2019, DOI: https://doi.org/10.18128/D080.V7.

21. Wakefield, J., Simpson, D. & Godwin, J. Comment: Getting into space with a weight problem. J Am Stat Assoc 111, 1111–1118,DOI: 10.1080/01621459.2016.1200918 (2016).

22. Diggle, P. & Ribiero Jr, P. Model-based geostatistics (Springer Series in Statistics, New York, 2007).

23. WorldPop: Population/Individual countries 2000-2020. https://www.worldpop.org/geodata/listing?id=29. Accessed 25 March 2020.

24. Version 4 DMSP-OLS Nighttime Lights Time Series. National Oceanographic and Atmospheric Administration. https://ngdc.noaa.gov/eog/dmsp/downloadV4composites.html. Accessed 30 March 2020.

25. IUCN: World Database on Protected Areas. https://www.iucn.org/theme/protected-areas/our-work/world-database-protected-areas. Accessed 09 Apr 2018.

26. Biodiversity A-Z. https://www.biodiversitya-z.org/content/protected-area. Accessed 01 Apr 2020.

27. World Wildlife Foundation. Global Lakes and Wetlands Database. https://www.worldwildlife.org/pages/global-lakes-and-wetlands-database. Accessed 09 Apr 2018.

28. U.S. Geological Survey. Digital Eelevation - Global Multi-resolution Terrain Elevation Data 2010 (GMTED2010). Accessed 01 Apr 2020.

29. Lindgren, F. & Rue, H. Bayesian spatial modelling with R-INLA. J. Stat. Softw. 63, 1–25 (2015).

30. Asmarian, N., Ayatollahi, S., Sharafi, Z. & Zare, N. Bayesian spatial joint model for disease mapping of zero-inflated data with R-INLA: A simulation study and an application to male breast cancer in Iran. Int J Environ Res Public Heal. 16, 4460, DOI: 10.3390/ijerph16224460 (2019).

31. Wakefield, J. Multi-level modeling, the ecologic fallacy, and hybrid study designs. Int. J. Epidemiol. 38, 330–336, DOI: doi:10.1093/ije/dyp179 (2009).

32. Fuglstad, G.-A., Simpson, D., Lindgren, F. & Rue, H. Constructing priors that penalize the complexity of Gaussian random fields. J Am Stat Assoc 114, 445–452, DOI: https://doi.org/10.1080/01621459.2017.1415907 (2019).

33. Simpson, D., Rue, H., Riebler, A., Martins, T. & Sorbye, S. Penalising model component complexity: A principled, practical approach to constructing priors. Stat Sci 32, 1–28, DOI: doi:10.1214/16-STS576 (2017).

34. Gilbert, M. et al. Global cattle distribution in 2010 (5 minutes of arc), DOI: 10.7910/DVN/GIVQ75 (2018).

35. Gilbert, M. et al. Global pigs distribution in 2010 (5 minutes of arc), DOI: 10.7910/DVN/33N0JG (2018).

36. Lumley, T. survey: analysis of complex survey samples (2016). R package version 3.32.

37. Mercer, L., Wakefield, J., Chen, C. & Lumley, T. A comparison of spatial smoothing methods for small area estimation with sampling weights. Spatial Stat. 8, 69–85 (2014).

